# Thyroid hormones mediate the impact of early-life stress on ventral tegmental area gene expression and behavior

**DOI:** 10.1101/2023.08.25.554785

**Authors:** Shannon N. Bennett, Austin B. Chang, Forrest D. Rogers, Parker Jones, Catherine Jensen Peña

**Author notes:** Correspondance to: Catherine J. Peña. These authors contributed equally to this work (Shannon N. Bennett, Austin B. Chang).

## Abstract

Proper thyroid function is essential to the developing brain, including dopamine neuron differentiation, growth, and maintenance. Stress across the lifespan impacts thyroid hormone signaling and anxiety disorders and depression have been associated with thyroid dysfunction (both hypo- and hyper-active). However, less is known about how stress during postnatal development impacts thyroid function and related brain development. Our previous work in mice demonstrated that early-life stress (ELS) transiently impinged on expression of a transcription factor in dopamine neurons shown to be regulated by thyroid hormones. We hypothesized that thyroid hormone signaling may link experience of ELS with transcriptional dysregulation within the dopaminergic midbrain, and ultimately behavior. Here, we find that ELS transiently increases thyroid-stimulating hormone levels (inversely related to thyroid signaling) in both male and female mice at P21, an effect which recovers by adolescence. We next tested whether transient treatment of ELS mice with synthetic thyroid hormone (levothyroxine, LT4) could ameliorate the impact of ELS on sensitivity to future stress, and on expression of genes related to dopamine neuron development and maintenance, thyroid signaling, and plasticity within the ventral tegmental area. Among male mice, but not females, juvenile LT4 treatment prevented hypersensitivity to adult stress. We also found that rescuing developmental deficits in thyroid hormone signaling after ELS restored levels of some genes altered directly by ELS, and prevented alterations in expression of other genes sensitive to the second hit of adult stress. These findings suggest that thyroid signaling mediates the deleterious impact of ELS on VTA development, and that temporary treatment of hypothyroidism after ELS may be sufficient to prevent future stress hypersensitivity.

## INTRODUCTION

Early life stress (ELS) alters brain development and is a major risk factor for anxiety, depression, and other psychiatric diseases (Bernet and Stein, 1999; Heim and Nemeroff, 2001; McLaughlin et al., 2010). While stress impacts many brain systems, reward circuitry is tightly linked to anhedonia, effort-based decision making, and cognitive behaviors often altered in mood disorders (Nestler and Carlezon, 2006; Russo and Nestler, 2013). Indeed, human and animal studies demonstrate long-lasting impact of ELS on reward system development, structure, function, and molecular regulation, which mediate altered behavioral differences (Duque-Quintero et al., 2022; Hanson et al., 2021). Recent work indicated that childhood adversity and socioeconomic disadvantage blunts development and connectivity of the ventral tegmental area (VTA) — the dopaminergic center of the brain’s reward circuitry — with target regions (Marusak et al., 2017; Park et al., 2021). ELS also alters dopamine release and metabolism, both at baseline and in response to stressors (Cabib et al., 1993; Matthews et al., 2001; Brake et al., 2004; Arborelius and Eklund, 2007). These structural and functional alterations are likely due to altered development at the molecular level (Parel and Peña, 2022). Indeed, ELS results in long-lasting transcriptional changes within VTA (Peña et al., 2017, 2019b). However, the molecular cascade of events linking experience of ELS with altered transcription in VTA is not fully understood.

We hypothesized that thyroid hormone signaling may be altered by ELS and upstream of transcriptional changes in VTA. Thyroid hormones (thyroxine, T4, and its active metabolite, triiodothyronine, T3) can induce dopamine neuron differentiation in cultured embryonic neurons (Lee et al., 2019). Hypothyroidism during embryonic development leads to a loss of dopamine neurons (Kincaid, 2001). These effects extend beyond the embryonic period, such that hypothyroidism in the neonatal period leads to dopaminergic dysfunction (Vaccari et al., 1990a; Vaccari et al., 1990b). The impact of thyroid hormone on dopamine neuron differentiation and maintenance is thought to be mediated by regulation of the transcription factor orthodenticle homeobox 2 (*Otx2*), an important determinant of dopamine neurogenesis and implicated in critical period plasticity (Sugiyama et al., 2008; Chen et al., 2015; Batista and Hensch, 2019). The two mammalian thyroid hormone receptors, THRα and THRβ, act as transcriptional activators when bound by ligand, and can directly regulate target genes and act as transcriptional repressors in the absence of ligand. Treatment of cultured neurons with T3 is sufficient to increase OTX2 protein levels, and blocking thyroid hormone with propylthiouracil (PTU) *in vivo* reduces OTX2 levels in VTA (Chen et al., 2015). *In vitro, Otx2* expression is critical for thyroid hormone-dependent dopamine neuron differentiation (Chen et al., 2015). Moreover, expression of *Otx2* is transiently suppressed in VTA by ELS, which mediates transcriptional and behavioral sequelae of ELS (Peña et al., 2017).

Stress across the lifespan has previously been shown to impact thyroid hormone signaling. Acute stress in adult rats and mice decreased peripheral thyroid hormone levels within two hours, an effect which lasted for up to twelve hours (Ahn et al., 2016; Helmreich and Tylee, 2011). Chronic stress in adulthood has also been found to alter thyroid hormone signaling in male rats, although both increases and decreases have been reported, which may depend on type and timing of stress exposure and sampling methodology (Guo et al., 2015; Kioukia-Fougia, 2002; Martí et al., 1993; Pollard et al., 1979). In rats, both maternal separation and low levels of maternal licking and grooming stimulation were associated with lower levels of T3 at different time points (Hellstrom et al., 2012; Jaimes-Hoy et al., 2021, 2016). In humans, post-traumatic stress disorder and depression have also been associated with thyroid dysfunction (both hypo- and hyper-active) (Fountoulakis et al., 2006; Jung et al., 2019; Wang and Mason, 1999). High levels of early life physical abuse in children have also been associated with lower levels of T3, after adjusting for pubertal status, sex, socioeconomic status, and BMI (Machado et al., 2015). However, the timing and duration of thyroid dysfunction following early-life stress is inconsistent across previous studies.

Here, we sought to understand whether and how long ELS dysregulates thyroid function, and whether acute pharmacological treatment can restore VTA gene expression profiles and behavioral sensitivity to stress in adulthood, in both male and female mice.

## METHODS

### Animals

All C57BL/6J mice were housed in a temperature-controlled environment and maintained on a 12 h light/ dark cycle (lights on at 04:00) with *ad libitum* access to food and water. All experiments involving animals were approved by the Institutional Animal Care and Use Committee at Princeton University and conducted in accordance with their guidelines. Mice were considered to be “male” or “female” based on external genitalia and/or anogenital distance (pups).

All experimental mice were generated from in-house breeding to avoid shipping stress during gestation. Virgin adult C57BL/6J mice were shipped from Jackson and mated in trios. Males were removed after 5 days, and females remained housed together until 1-3 days prior to giving birth, at which point they were given individual cages. Litters were randomly assigned at birth to standard-rearing (Std) or early life stress (ELS) conditions, as described below. All offspring were weaned at postnatal day P21 by sex. All experiments included both male and female offspring. One cohort was generated for the hormone assay time-course, and a second cohort was used for pharmacological treatment and behavior and gene expression experiments.

### Early life stress paradigm

ELS occurred from P10-17 as previously described (Peña et al., 2017, 2019a, 2019b; Balouek et al., 2023; Parel et al., 2023) and consisted of both maternal separations (with all pups from a litter removed to a clean cage and returned to the home cage 3-4 hours later) and limited nesting material in the home cage (to ½ standard EnviroDri puck size). Pups were given access to diet gel and several chow pellets on the cage floor during separations, although pups did not appear to eat during separation. As an additional stressor, soiled bedding from retired Swiss-Webster male aggressor mice was added daily to the separation cages of pups during separation window. At the conclusion of ELS on P17, mice were given clean cages with standard cage enrichment, consisting of a hut and full puck of nesting material, and left undisturbed until weaning.

### Thyroid-Stimulating Hormone (TSH) Assay

Trunk blood was taken from male and female Std and ELS mice at three ages: P21, P40, and P60 (N=5-9 per sex per group). Blood was collected into sterile tubes coated with 1.5M EDTA. Tubes were placed on wet ice and centrifuged at 3000 G at 4ºC for 15 minutes. Resulting supernatant plasma was collected and transferred to new sterile tubes for storage at -80ºC until use. An enzyme-linked immunosorbent assay (ELISA) kit (#RK03260, Abclonal Technology, Woburn, MA) was used to assess plasma TSH levels. The stated assay sensitivity is 19.2 pg/mL. ELISAs were performed with the provided standards according to the manufacturer’s protocol. Plates were read on either on a Spectramax iD5 (P21) or a Biotek Gen 5 (P40 & P60) plate readers, at 450 nm wavelength was used for analysis. All standards and samples were run in duplicate with high correlation among replicates (R^2^=0.9373). Samples from different ages were processed on different days.

### Synthetic thyroid hormone treatment (LT4)

In a second cohort, litters were randomly assigned to one of three rearing conditions: Std, ELS, or ELS+LT4. ELS was performed identically for ELS and ELS+LT4 groups. Levothyroxine (LT4, synthetic T4 hormone), which is then converted to active T3 in the body, is the gold standard in treating hypothyroidism (Garber et al., 2012). All animals in this cohort were introduced to water bottles filled with tap water at P17 as training, while normal lixit water spouts were removed. Upon weaning at P21, mice in the ELS+LT4 group were given 1.0 μg/mL LT4 in drinking water for an approximate consumption of 0.2 - 0.3 mg/kg/ day, based on 2-3 mL average daily water consumption at P21. Supplementation continued for four additional days until P25. For each cage, one 0.1 mg tablet of LT4 (Henry Schein, Baltimore MD, #1381054) was first dissolved in 5 mL of drinking water with 1 mM (1 μL 5M) NaOH, then topped up to 100 mL of drinking water and pH’d to neutral as needed. Treatment dosage and duration was based on previous findings that 1.0 μg/mL could effectively elevate thyroid hormone levels within one week, while 3.0+ μg/mL of T3 was shown to be thyrotoxic (Davenport et al., 1975; East-Palmer et al., 1995; Engels et al., 2016; Niedowicz et al., 2021), and we sought only to rescue thyroid to endogenous levels rather than induce hyperthyroidism.

### Adult chronic non-discriminatory social defeat stress (CNSDS)

In order to assess the impact of early rearing and LT4 treatment conditions on stress susceptibility in adulthood, male and female mice from each group were randomly assigned to control or chronic non-discriminatory social defeat stress (CNSDS) groups, performed as previously described (Golden et al., 2011; Yohn et al., 2019). Briefly, beginning at ages P68-P70, CNSDS-assigned mice were subject to 10 days of daily social defeat by a Swiss Webster retired breeder (hereafter, “aggressor;” Taconic) in a standard rat cage filled with corn cob bedding. An experimental male mouse was introduced to an aggressor’s cage first for 3 minutes, followed by an experimental female mouse for an additional 5 minutes. Males were then moved across a perforated plexiglass barrier within the aggressor cage for the remainder of the day, for sensory stress without physical interaction. Females were removed to individual cages with aggressor bedding for the remainder of the day. Experimental mice were introduced to a new aggressor each day for 10 days, and males and females rotated in different directions so that trios were unique each day. While aggressors were observed to attack females, attacks were predominantly directed towards males. No copulations were observed. Control mice were housed in a standard mouse cage in pairs of the same sex, separated by a perforated plexiglass barrier, and also rotated. At the conclusion of CNSDS, male and female mice were rehoused in clean cages and underwent two consecutive days of behavioral testing.

### Behavior

#### Social Avoidance Testing

On the day following CNSDS, experimental mice were tested in a two-stage social avoidance test under red lighting, as previously described (Berton et al., 2006; Peña et al., 2017). Social avoidance has been previously associated with other depression-like behaviors and is responsive to antidepressant treatment (Berton et al., 2006; Krishnan et al., 2007). In the first 2.5-min stage, the experimental mouse was allowed to freely explore an arena (44×44x20 cm) containing a plexiglass and wire mesh enclosure (novel object; 10×6 cm) centered against one wall of the arena. In the second 2.5 min stage, the experimental mouse was immediately returned to the arena with a novel Swiss Webster mouse (aggressor strain) enclosed in the plexiglass wire mesh cage. Time spent in the ‘interaction zone’ (14×26 cm) surrounding the plexiglass wire mesh cage, ‘corner zones’ (10×10 cm), and ‘distance traveled’ within the arena was measured by video tracking software (Ethovision, Noldus). A social interaction ratio (SI Ratio) was calculated of time spent exploring the novel mouse over time exploring the novel object; mice were considered “susceptible” to CNSDS if SI Ratio<0.9, “resilient” if SI Ratio >1.1, and “indifferent” for interaction scores in between. All males were tested before females. Within-sex, testing order was counterbalanced by group.

#### Open Field Testing

On the second day of behavioral testing, exploration of an open field arena (44×44 cm) was assessed during a 10 min test under red lighting, as previously described (Peña et al., 2017). A video-tracking system (Ethovision, Noldus) measured locomotor activity, as well as the time spent in the center (34×34 cm) and periphery of the test arena as an index of anxiety. Mice were considered “susceptible” to CNSDS if Open Field Center Time was less than 1 standard-deviation from the mean of control mice of the same sex, and otherwise “resilient.”

### Real-time qPCR

On the day following behavioral testing, mice were sacrificed by rapid cervical dislocation. Brains were rapidly removed, and bilateral 1 mm-thick 16 gauge punches of VTA were taken fresh, flash-frozen on dry ice, and stored at -80C until processing. RNA was extracted from tissue samples with Trizol (Invitrogen) and chloroform (Sigma) and purified with RNeasy Micro Kits(Qiagen). cDNA was created with are verse-transcription kit (HighCapRT, Applied Biosystems). Primers for real-time semi-quantitative PCR (qPCR) were validated for efficiency and specificity (Table 1). All qPCR reactions were run in triplicate and used SYBR-green on a QuantStudio 7 (ThermoFisher). The 2^(-ΔΔCt) method was used to calculate RNA expression analysis compared to the standard-reared group (including both males and females) with Hprt as a control reference gene (Livak and Schmittgen, 2001).

**Table 1:**
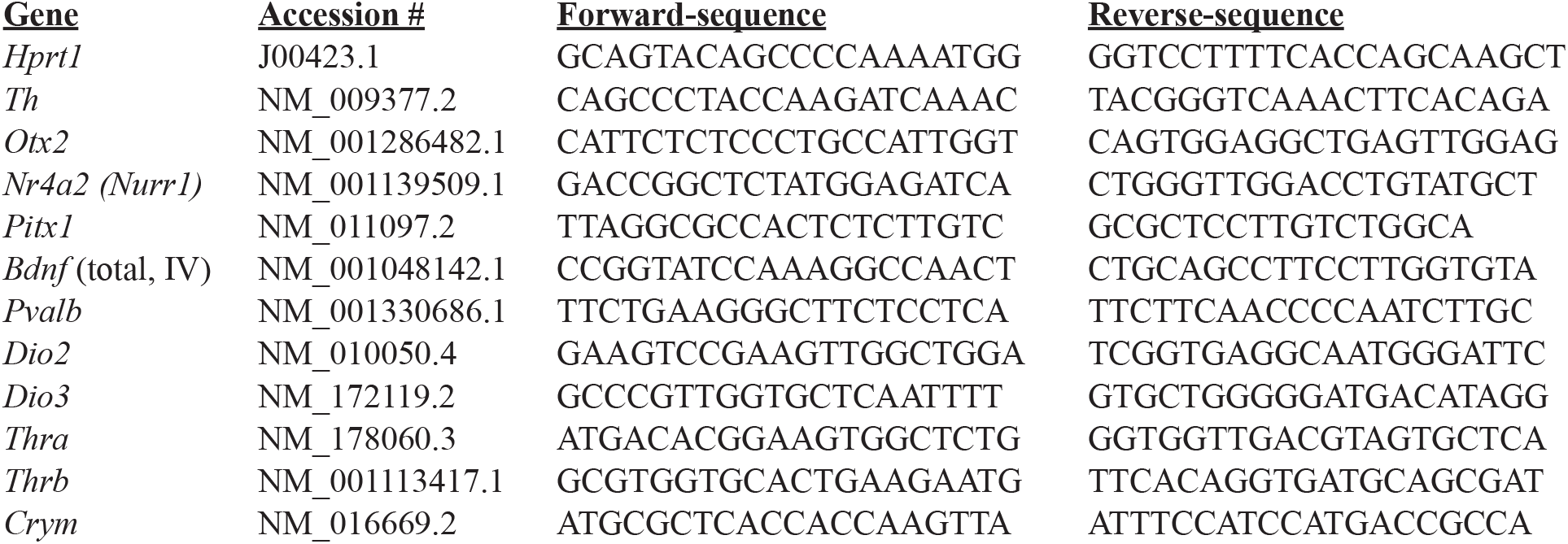
Primers for qPCR.

### Statistical Analysis

All statistics were performed using SPSS (Version 26) or Prism (GraphPad; version 9). Outliers were defined as more than 2 standard deviations from the group mean and were removed prior to inferential analyses. Main effects and interactions were determined using general linear model (examining interactions between sex, early rearing group, and CNSDS) or ANOVA (effect of early rearing group only) with Tukey’s HSD post hoc analysis. Effect sizes for ANOVA were calculated and reported as eta2. Differences in proportions were tested using Kruskal-Wallis non-parametric test, run on resilience membership only. All significance thresholds were set at *p* ≤ 0.05.

## RESULTS

### Early life stress transiently increases thyroid-stimulating hormone

In order to determine whether and how long ELS disrupts thyroid hormone signaling, we examined thyroid-stimulating hormone (TSH) in males and females at three time points (**Figure 1A**): P21 prior to weaning, P40 during adolescence, and P60 in young adulthood. TSH was used as a proxy for thyroid hormone function, as TSH is inversely related to thyroid hormone levels: TSH promotes synthesis and release of of T4 and T3 from the thyroid gland, and is negatively regulated by increasing levels of thyroid hormones in circulation (Feldt-Rasmussen et al., 2021; Yen, 2001). At P21, there was a main effect of sex [F(1, 19)=14.36, *p*=0.008, η^2^=0.21] and a main effect of ELS [F(1, 19)=8.786, *p*=0.001, η^2^=0.57], such that female mice had higher levels of TSH and ELS increased TSH among both males and females (**Figure 1B**). The effect of ELS on TSH recovered by adolescence and was no longer significant (*p*>0.20 for all main effects and interactions at P40 and P60; **Figure 1C-D**).

**Figure 1.**
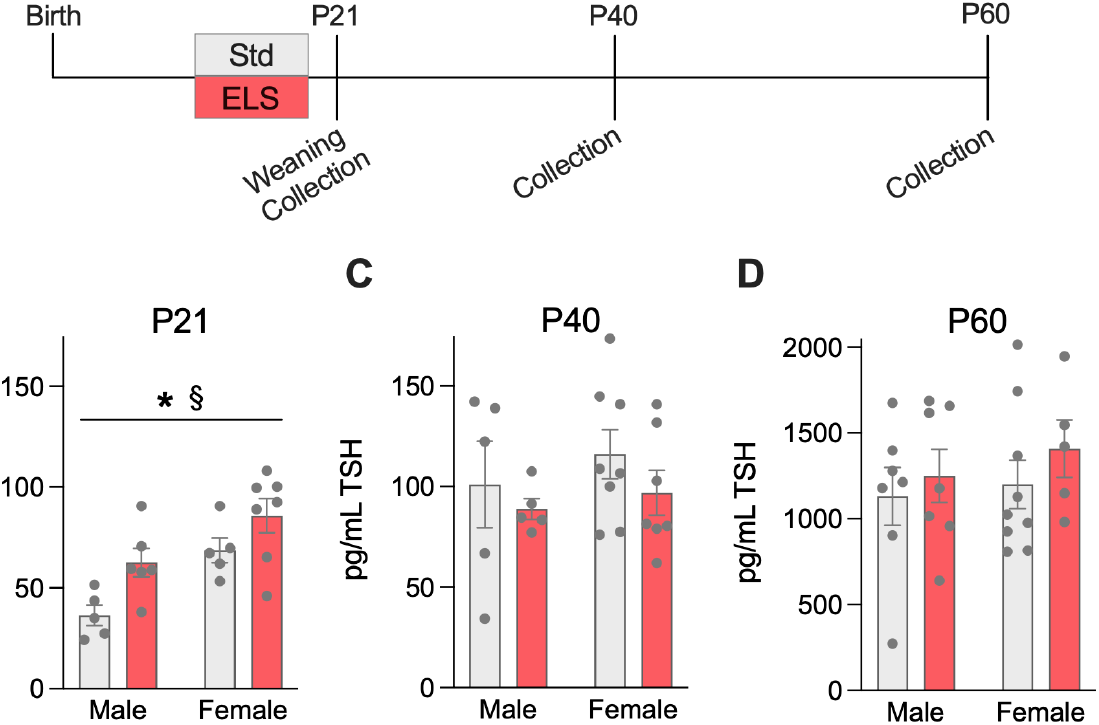
ELS increases thyroid-stimulating hormone levels at P21. Experimental timeline (**A**). Plasma TSH levels at P21 (**B**), P40 (**C**), and P60 (**D**). Error bars indicate mean ± SEM. * main effect of ELS (*p*<0.05); § main effect of sex (*p*<0.05).

### LT4 supplementation following early life stress rescues adult-stress-induced behavioral deficits in male mice

In order to determine whether temporary juvenile thyroid rescue could mitigate the impact of ELS on behavioral stress sensitivity previously described (Peña et al., 2017, 2019a, 2019b; Balouek et al., 2023), we tested response to chronic stress in adulthood on two different behavioral tasks of social avoidance behavior and exploration of a novel arena (**Figure 2A**). Stressed mice underwent chronic non-discriminatory social defeat stress (CNSDS), a form of social stress that can be simultaneously used with both male and female mice (Yohn et al., 2019). We found a main effect of sex [F(1,51)=35.375, *p*<0.001, η^2^=0.10], a main effect of early life rearing group [F(2,51)=6.359, *p*=0.003, η^2^=0.014], and a main effect of CNSDS [F(1,51)=4.842, *p*=0.032, η^2^=0.002] on social interaction time. Given the main effect of sex and our prior work showing no effect of ELS alone, we examined the impact of early-life condition within control and CNSDS males and females separately. We found an effect of early-life condition on social interaction time within CNSDS males [F(2,10)=8.213, *p*=0.012, η^2^=0.057; **Figure 2B**]. Post-hoc analysis with Tukey’s HSD multiple comparisons correction confirmed that ELS reduced social interaction time compared to Std (*p*=0.033), an effect which was lost with LT4 treatment (*p*=0.298). The effects of ELS and LT4 treatment were not found among females (*p*=0.196; **Figure 2C**), nor among control mice (*p*>0.38). We also examined the proportions of mice considered resilient, indifferent, or susceptible to stress following CNSDS (see methods), and found that ELS increased susceptibility (particularly among males similar to previous reports; (Peña et al., 2017, 2019a)), and that treatment with LT4 after ELS restored resilience [among CNSDS males: *H*(2)=6.333, *p*=0.042; among CNSDS females: *H*(2)=5.0, *p*=0.082; **Figure 2D**]. In the open field test, there was a main effect of sex [F(1,54)=6.306, *p*=0.015, η^2^=0.005] and a main effect of CNSDS [F(1,54)=11.137, *p*=0.002, η^2^=0.016] on time spent exploring the center of the open arena (**Figure 2 E-F)**. Among males, there was a main effect of CNSDS to reduce center exploration [F(1,25)=8.772, *p*=0.007, η^2^=0.123] with a small effect of early rearing condition [F(2,25)=2.047, *p*=0.150, η^2^=0.027]. Among males, but not females, treatment with LT4 after ELS appeared to increase the proportion of mice resilient to CNSDS on this behavior [among CNSDS males: *H*(2)=8.1, *p*=0.017; **Figure 2G**].

**Figure 2.**
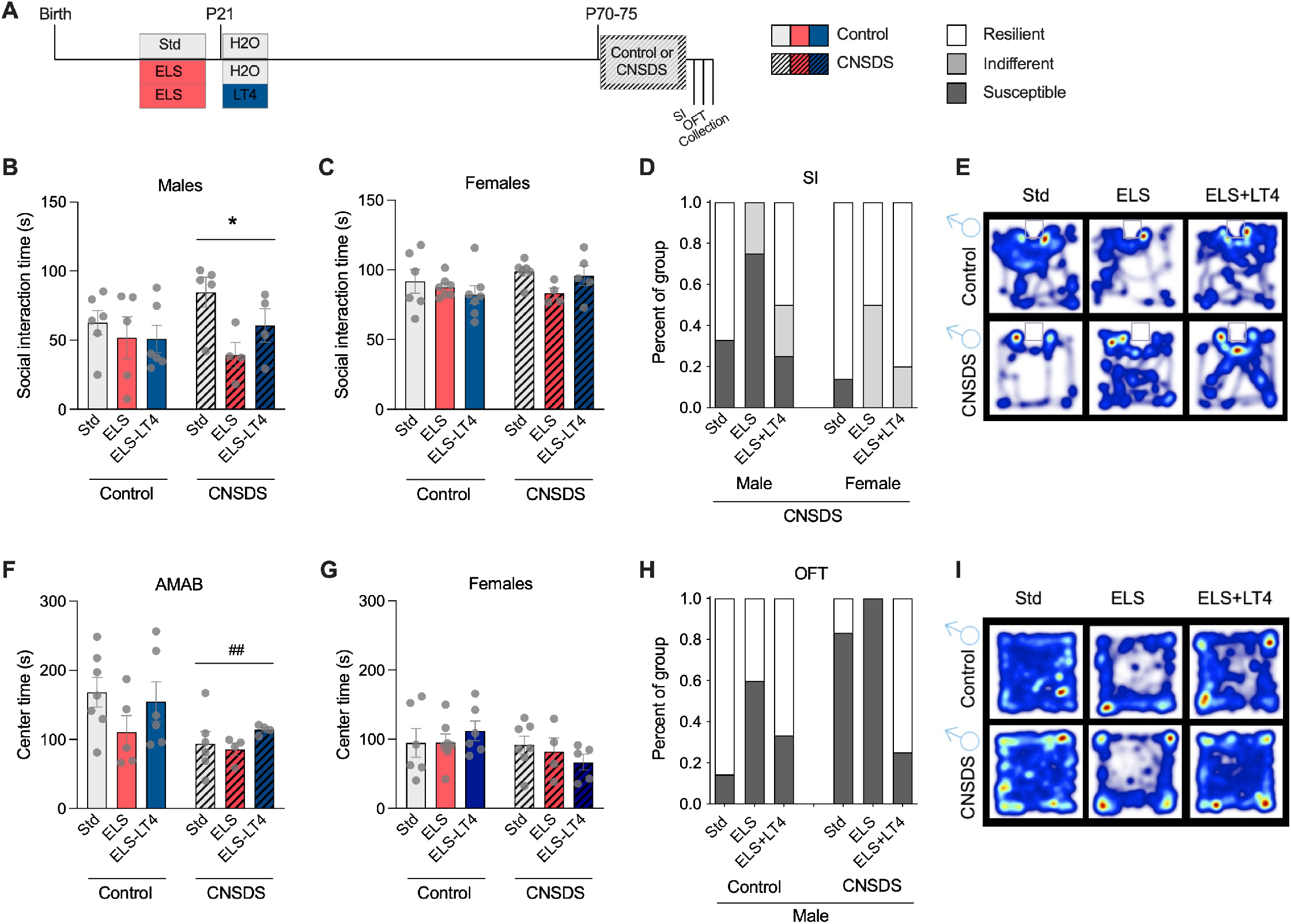
Juvenile LT4 treatment rescues stress-induced behavioral changes in males. Experimental timeline and groups (**A**). Mice were given either standard drinking water or water supplemented with LT4 from P21-P25. Time spent investigating a novel aggressive mouse among males (**B**) and females (**C**). Among mice that underwent chronic non-discriminatory social defeat stress, percent of each group considered resilient, indifferent, or susceptible on the social interaction test (**D**). Representative heatmaps of mouse exploration during social interaction testing among males, with novel social targets enclosed at the top center of each arena; warmer colors represent more time at that location (**E**). Time spent exploring the center of an open field among males (**F**) and females (**G**). Among male mice, percent of each group considered resilient or susceptible on the open field test (**H**). Representative heatmaps of mouse exploration during open field testing among males (**I**). Error bars indicate mean ± SEM. * Main effect of early life condition (*p*<0.05). ## Main effect of CNSDS (*p*<0.01).

### LT4 supplementation following early life stress rescues expression of thyroid and dopamine related gene expression in the VTA

Thyroid signaling has been implicated as an upstream regulator of dopamine neuron development in VTA (Chen et al., 2015; Crocker and Overstreet, 1983; Kreutz et al., 1990; Vaccari et al., 1990a; Vaccari et al., 1990b), and we previously demonstrated the importance of transcriptional regulation in VTA for ELS-induced behavior changes (Peña et al., 2017, 2019b; Kronman et al., 2021). In order to determine whether temporary juvenile thyroid rescue could mitigate the impact of ELS on transcription within VTA, we assessed expression of genes related to VTA maturation and function (tyrosine hydroxylase, *Th*; orthodenticle homeobox 2, *Otx2*; *Nurr1* or *Nr4a2*, and *Pitx1;* **Figure 3B-E**), plasticity (brain derived neurotrophic factor, *Bdnf*; and parvalbumin, *Pvalb*; **Figure 3F and L**) and local thyroid function (the deiodinases *Dio2* and *Dio3*, the thyroid hormone receptors *Thra* and *Thrb*, and the thyroid hormone binding protein Crystalin mu, *Crym*; **Figure 3G-K**) by qPCR. We found a main effect of early life condition on expression of *Pvalb* [F(2,37)=5.761, *p*=0.007, η^2^=0.044], *Thra* [F(2,37)=5.127, *p*=0.011, η^2^=0.017], and *Thrb* [F(2,37)=5.371, *p*=0.009, η^2^=0.042], with trends among *Nr4a2* [F(2,37)=2.714, *p*=0.079, η^2^=0.008] and *Crym* [F(2,37)=2.997, *p*=0.062, η^2^=0.005]. There was a main effect of CNSDS on expression of *Th* [F(1,37)=27.701, *p*<0.001, η^2^=0.133], *Nr4a2* [F(1,37)=7.33, *p*=0.010, η^2^=0.014], and *Dio2* [F(1,37)=6.791, *p*=0.013, η^2^=0.017]. There were no main effects of sex. There were significant interactions between sex and early life condition for expression of *Otx2* [F(2,37)=3.844, *p*=0.030, η^2^=0.021], *Nr4a2* [F(2,37)=6.182, *p*=0.005, η^2^=0.041], *Dio3* [F(2,37)=3.489, *p*=0.041, η^2^=0.015], *Thra* [F(2,37)=5.074, *p*=0.011, η^2^=0.017], and *Crym* [F(2,37)=3.426, *p*=0.043, η^2^=0.006]. There was also an interaction between sex, early life condition, and CNSDS on expression of *Crym* [F(2,37)=10.768, *p*<0.001, η^2^=0.061] and *Thra* [F(2,37)=5.056, *p*=0.011, η^2^=0.017], with trends for expression of *Bdnf* [F(2,37)=3.002, *p*=0.062, η^2^=0.014] and *Dio3* [F(2,37)=2.477, *p*=0.098, η^2^=0.008].

**Figure 3.**
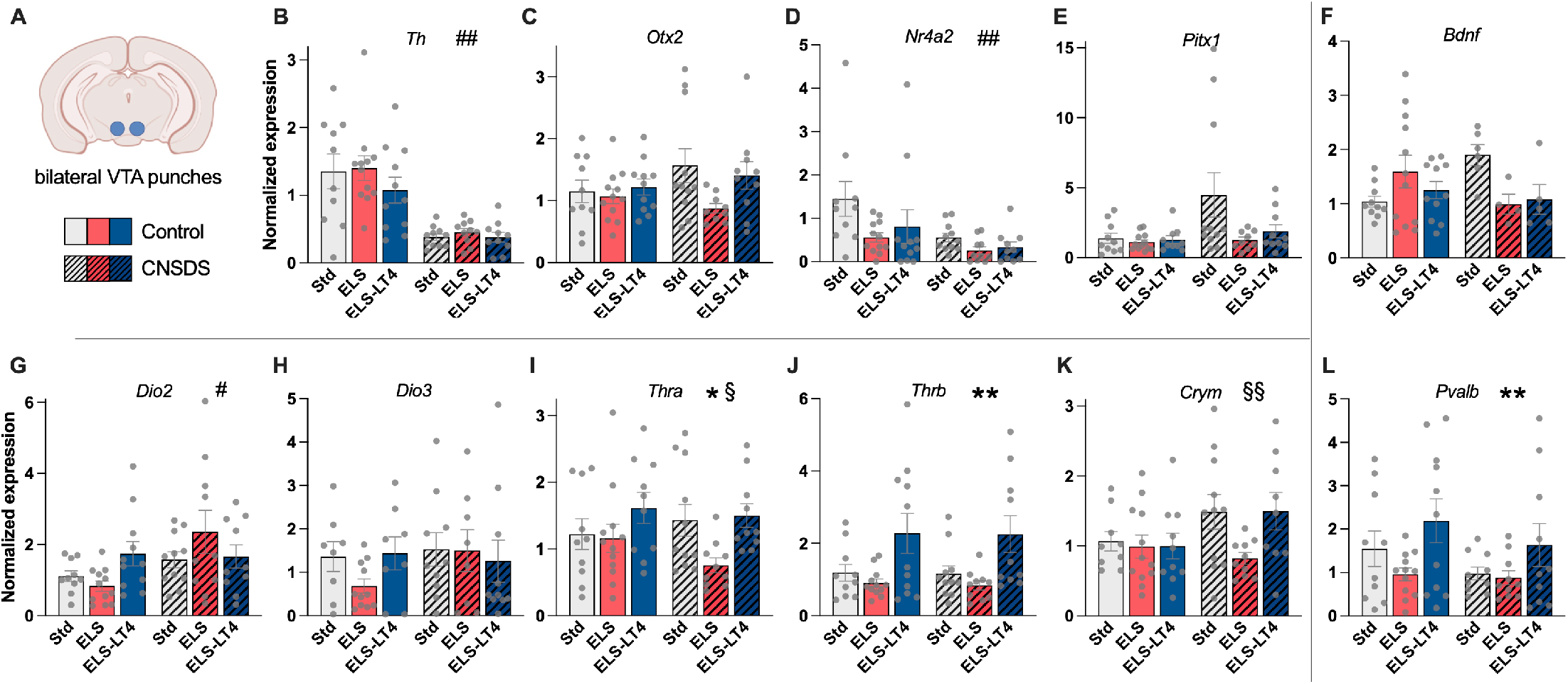
LT4 treatment ameliorates stress-induced changes in gene expression in adult VTA. Bilateral tissue punches were taken of adult VTA one day after completion of behavioral testing (A). Relative expression was measured by qPCR for genes related to VTA maturation including tyrosine hydroxylase (*Th*; B), orthodenticle homeobox 2 (*Otx2*; C), *Nr4a2* (also known as *Nurr1*; D), and *Pitx1* (E); genes related to plasticity including brain-derived neurotrophic factor (*Bdnf*; F), and parvalbumin *(Pvalb;* L); and genes related to thyroid hormone processing and signaling including deiodinase 2 (*Dio2*; G), *Dio3* (H), thyroid hormone receptor-alpha (*Thra*; I), thyroid hormone receptor-beta (*Thrb*; J), and crystallin mu (*Crym*; K), * Main effect of early life condition. Error bars indicate mean ± SEM. # Main effect of CNSDS. § Interaction between early life condition, CNSDS, and sex.

## DISCUSSION

Thyroid hormone signaling is essential for brain development and has been historically linked with depression, anxiety, and cognition, but there is a critical gap in our understanding of how stress during development may impinge on thyroid function to mediate mood disorder risk. We sought to link these elements. Our prior work demonstrated that ELS increases stress sensitivity via transcriptional changes in VTA mediated by transient developmental disruption of the transcription factor OTX2 (Peña et al., 2017). Moreover, there is *in vitro* evidence that thyroid hormone acts upstream of OTX2 to regulate dopamine neuron maturation (Chen et al., 2015). We therefore specifically sought to understand whether altered thyroid function mediates the impact of ELS on factors essential for VTA development, and ultimately behavioral stress sensitivity in male and female mice. We found that ELS transiently increased TSH, but recovered by adolescence. We also found evidence that short-term treatment with synthetic thyroid hormone following ELS rescues aspects of ELS-induced sensitivity to adult stress on VTA gene expression and behavior, particularly among male mice. These data provide evidence that altered thyroid hormone signaling may link experience of ELS with downstream maturation of dopaminergic brain regions and behavioral stress sensitivity.

### ELS transiently suppresses thyroid function

We found that ELS transiently increased plasma TSH levels in juvenile male and female mice at P21, indicative of suppressed T3 and T4 levels and impaired thyroid function for at least four days following ELS. In our mouse model of ELS, however, TSH levels recovered by adolescence. Prior research indicated that stress alters thyroid function, although the direction and duration of effect is inconsistent in the literature. For example, maternal separation from P2-21 in rats was previously found to have no impact on TSH among adolescent (P40) males or females but to decrease both TSH and T3 in adulthood (P90) (Jaimes-Hoy et al., 2016), and to increase TSH among adult (P80) males in another study (Jaimes-Hoy et al., 2021). Adult rats subject to two weeks of repeated stress also had significantly reduced total and free T3 and T4, measured the day after stress ended (Helmreich et al., 2005; Zhang et al., 2018). Another study found that one day of stress in adult male rats was insufficient to alter thyroid hormone levels, but that T3 and T4 were significantly reduced by at least three days of stress (Servatius et al., 2000). Among humans, high levels of childhood trauma were associated with lower plasma T3 (but not T4), measured in adolescence (Machado et al., 2015). Women with a history of child abuse were also found to be at higher risk for thyroid dysfunction later in life, particularly during periods of endocrine change such as pregnancy and postpartum (Moog et al., 2017; Plaza et al., 2010). Thus across studies and age of stress, chronic stress appears to impinge on thyroid function at least temporarily. The effect of at least temporarily suppressed thyroid function is likely to have a larger impact among children whose brains and bodies are still developing and for whom thyroid dysfunction may alter entire developmental trajectories, compared to adults for whom only steady-state processes may be temporarily impacted. For example, our previous work demonstrated that even temporary suppression of the transcription factor *Otx2* during postnatal development — but not adulthood — had long-lasting consequences on downstream cascades of gene expression and behavior (Peña et al., 2017). Together with previous studies, our findings of the impact of ELS on thyroid function call for children and adolescents who have experienced chronic stress or trauma to be screened for thyroid dysfunction.

### Juvenile LT4 treatment rescues stress-induced behavioral changes in males

We hypothesized that if thyroid dysfunction mediated the impact of ELS on hypersensitivity to later stress, rescue of thyroid signaling after ELS with levothyroxine treatment would ameliorate ELS-induced behavioral changes. Because mice showed short-term impairment of thyroid function after ELS that recovered by adolescence, and the half-life of LT4 is approximately 7 days (Lipp, 2021), we treated mice for only five days with LT4 in drinking water in order to prevent over-treatment. Consistent with previous findings (Peña et al., 2017, 2019b), the combination of ELS and adult social defeat stress increased social avoidance in male mice, and to a lesser extent in female mice (Figure 2B-E). Here, we found that juvenile LT4 treatment prevented ELS-induced hypersensitivity to adult stress and instead promoted resilience. Increased anxiety- and depression-like behavior is consistent with a hypothyroidic state in adult rodents (Wilcoxon et al., 2007; Buras et al., 2014; Niedowicz et al., 2021), despite our findings of normal TSH levels in adulthood, further indicating the potential long-term effects of juvenile thyroid disruption. However, it is possible that the second hit of stress in adulthood itself disrupted thyroid function, as indicated by a previous study that observed hypothyroidism in socially defeated male rats (Olivares et al., 2012).

The impact of adult social defeat stress was milder among female mice. Similarly, our recent work showed that female mice remained social after social defeat stress and that novelty-suppressed feeding may be a better behavioral readout of stress for females (Parel et al., 2023). Here, we sought to compare sexes directly and therefore use the same behavioral tests for all animals, and we chose not to subject mice to food restriction for novelty-suppressed feeding testing given the role of thyroid hormones in metabolism and potential interactions on downstream gene expression. Particularly given the main effect of sex on TSH levels and higher prevalence of depression among women, future studies should explore a wider variety of chronic adult stressors and behavioral measures that may be more ethological or better tailored for use with female rodents (Lopez and Bagot, 2021).

### LT4 treatment ameliorates stress-induced changes in gene expression in adult VTA

A growing body of work implicates VTA function in depression and anxiety-related behaviors (Berton et al., 2006; Chaudhury et al., 2013; Friedman et al., 2014; Hanson et al., 2021; Krishnan et al., 2007; Tye et al., 2013; Willmore et al., 2023). Our previous work found ELS led to enduring changes in gene expression in VTA, mediated by transient suppression of the transcription factor *Otx2* (Peña et al., 2017). Research *in vitro* indicated that OTX2’s role in maturation of dopamine neurons was downstream of thyroid hormone (Chen et al., 2015). We therefore hypothesized that thyroid hormone signaling could represent a missing link between stress experience and *Otx2*-mediated changes in dopamine neuron development in VTA. We profiled genes related to dopamine neuron development and function (*Th, Otx2, Nr4a2, Pitx1*), plasticity (*Bdnf, Pv*), and thyroid signaling (*Dio2, Dio3, Thra, Thrb*, and *Crym*). Among a subset of genes, juvenile LT4 treatment appeared to rescue the impact of ELS on expression prior to adult stress, including *Pvalb, Bdnf, Nr4a2*, and *Dio3*. Among other genes, juvenile LT4 treatment appeared to prevent abnormal response to adult stress, including *Thra, Crym*, and *Otx2. Crym* (crystallin mu, a thyroid hormone binding protein) has been previously implicated in sex-specific effects of social stress during development (Walker et al., 2022). The thyroid receptors THRα and THRβ act as transcriptional repressors in the absence of ligand, and transcriptional activators in the presence of thyroid hormone (Hörlein et al., 1995). Here, we see that two hits of stress reduces *Thra* which would predict blunted gene expression response to adult stress. We see this in a failure to express several genes normally upregulated by adult stress, including *Bdnf, Otx2*, and *Crym*. Rescuing developmental deficits in thyroid hormones after ELS not only restores levels of *Nurr1, Bdnf, Dio3*, and *Pvalb* after ELS alone, but also maintains normal levels of *Thra* and elevates *Thrb*, which may facilitate expression of downstream genes including *Otx2*, restoring levels of which has already been demonstrated to rescue depression-like behavior in the face of adult social stress. These results therefore provide initial evidence that rescuing thyroid hormone signaling after ELS rescues changes in dopamine neuron development in VTA. However, additional work needs to be done to determine the sufficiency of impaired thyroid signaling within VTA on these factors, and determine the impact of thyroid rescue within other stress-sensitive brain regions.

### Altered thyroid hormone function is implicated in depression, anxiety disorders, and cognition

Hypothyroidism and elevated TSH levels have been linked with neuropsychiatric disorders including depression (Bauer et al., 2008; Forman-Hoffman and Philibert, 2006; Saxena et al., 2000). While abnormalities in thyroid function have been reported among depressed patients (Kafle et al., 2020; Radhakrishnan et al., 2013), the direction of causality has been debated (Marangell and Callahan, 1998; Musselman and Nemeroff, 1996). Interestingly, treatment of euthyroid patients (patients within a normal range of thyroid hormone levels) with synthetic thyroid hormone has been shown to increase efficacy and accelerate response of antidepressant treatment, further bolstering the link between thyroid signaling and mood and anxiety disorders (Abraham et al., 2006; Agid and Lerer, 2003; Bauer, 1998; Bauer and Whybrow, 2021; Joffe et al., 1995; Rosenthal et al., 2011). The mechanisms of these effects are poorly understood, although metabolic effects and rescued activity in limbic brain regions with LT4 treatment has been reported (Bauer and Whybrow, 2021). Additional research is needed to explore the intriguing possibility that depressed patients with a history of ELS have forms of depression that may specifically benefit from thyroid hormone-augmented antidepressant treatment, even in adulthood when thyroid levels may be within normal range.

## Conclusions

Our findings suggest the detrimental effects of ELS on VTA development are mediated by thyroid hormone signaling and that short-term treatment with synthetic thyroid hormone following ELS may prevent susceptibility to future stress. Together with other studies, our work suggests that child and adolescent populations that have experienced chronic stress and trauma should screen for thyroid dysfunction. Additional pre-clinical and clinical research is needed to determine whether different forms of childhood stress and trauma impair thyroid function, the duration of such changes, and other brain systems that may be impacted. Ultimately, the treatability of thyroid dysfunction provides an attractive potential target for preventing or ameliorating the long-term impact of ELS on psychiatric disease susceptibility.

## ACKNOWLEDGEMENTS

This research was funded by NIH R00MH115096 (CJP); NIH R01MH129643 (CJP); PNI Research Innovator Award (CJP); New York Stem Cell Foundation (CJP); Princeton Office of Undergraduate Research (ABC). CJP is a New York Stem Cell Foundation Robertson Investigator and Robin Chemers Neustein Fellow. We thank Dr. Deena Walker for her conversations and insights related to this research.

## AUTHOR CONTRIBUTIONS

CJP and SNB designed the studies. SNB, ABC, FDR, and PJ collected the data. SNB, ABC, and CJP analyzed the data and created figures. SNB and CJP wrote the manuscript with input from all authors. All authors approve the manuscript.

